# Mapping memory-guided saccade population receptive fields reveals principles of cortical organization along the human intraparietal sulcus

**DOI:** 10.64898/2026.01.13.699231

**Authors:** Alessio Fracasso, Jasper Fabius, Katarina Moravkova, Rosanne Timmerman, Rebecca Taylor, Antimo Buonocore

**Author notes:** **Corresponding author** Alessio Fracasso. Statements and Declarations: **Nothing to disclose.**.

## Abstract

The intraparietal sulcus is a central hub for motor planning and sensorimotor integration. Its responses are known to support coordinate transformations across reference frames and modalities—for example, eye-position gain fields that convert retinotopic, fovea-centred signals into head-centred representations.

In the oculomotor domain, the human intraparietal sulcus is sensitive to eye-movement planning and has been extensively studied using memory-guided saccade paradigms. These classic approaches have revealed topographic organization and contralateral sensitivity for preferred direction in saccade planning and execution. However, such designs provide detailed information about preferred direction; they do not probe tuning width: how narrowly or broadly a given cortical location responds around its preferred direction.

At present, it remains unclear whether and how tuning width is organized along the human intraparietal sulcus. Critically, neurophysiological studies in non-human primates do not offer systematic and extensive measurements of tuning width. Moreover, the relationship between tuning width and preferred direction at the population level in human intraparietal cortex is unknown.

Here, we address these questions using a forward-modelling approach combined with high-field (7-Tesla) functional MRI. We observe contralateral preference for preferred direction in saccade planning and execution, as well as a novel large-scale organization of preferred saccade tuning width, in vivo, in humans.

**Highlights:**

saccade planning and execution tuning width is arranged topographically along human early visual cortex and human intraparietal sulcus;
specificity of voxel-level saccade planning and execution decreases along the visual hierarchy;
specificity of population-level saccade planning and execution increases along the visual hierarchy;
forward modelling applied to memory-guided saccade paradigm outperforms the standard phase-encoding approach;

## Introduction

Studies in humans and non-human primates have established the intraparietal sulcus (IPS) as a central hub for motor planning and sensorimotor integration (Mountcastle et al., 1975; Robinson et al., 1978; Corbetta and Shulman, 2002; Swisher et al., 2007). Cortical regions within human IPS are sensitive to eye-movement planning and execution, as well as to stimulus motion (Kastner et al., 2007; Konen and Kastner, 2008; Silver and Kastner, 2009). However, IPS responses are not limited to attentional or sensorimotor processing: its responses also correlate with higher-order cognitive functions, including decision-making (Shadlen and Newsome, 1996; Platt and Glimcher, 1999; Roitman and Shadlen, 2002; Glimcher, 2003; Dorris and Glimcher, 2004), as well as numerosity and quantity estimation (Harvey et al., 2013; Harvey et al., 2015; Harvey et al., 2020; van Dijk et al., 2021).

IPS activity has additionally been shown to support coordinate transformations across reference frames and modalities—for example, through eye-position gain fields that facilitate the conversion of retinotopic, fovea-centred signals into head-centred representations (Andersen et al., 1985; Duhamel et al., 1997; Bremmer et al., 2002; Levy et al., 2007; Xu et al., 2012; Konen et al., 2013; Merriam et al., 2013; Fabius et al., 2016; Fabius et al., 2019; Fabius et al., 2020; Fabius et al., 2022). Together, this evidence highlights the multimodal nature of IPS responses and their involvement across multiple effectors and behavioural contexts.

In the oculomotor domain, human IPS sensitivity to eye-movement planning and execution has been repeatedly demonstrated using the classic memory guided saccade task adopted by Sereno and colleagues (Sereno et al., 2001), typically implemented through phase-encoded (travelling-wave) designs. This analytical approach has been successfully adopted by several studies to obtain estimates of sensitivity across different parameter spaces in human cortex and subcortex (Engel et al., 1994; Engel et al., 1997; Schneider et al., 2004; Dumoulin et al., 2017b; Huang et al., 2017; Fracasso et al., 2021).

In phase-encoded designs involving eye movements, participants make saccades to remembered peripheral targets arranged cyclically around the visual field, following either clockwise or counter clockwise sequences. The resulting responses are analysed by fitting a sine wave to extract the optimal phase of the signal, which is lawfully related to saccade direction. This approach yields robust estimates of preferred orientation (tuning location) for each voxel, and is computationally efficient (Sereno et al., 2001; Konen et al., 2004; Schluppeck et al., 2005; Kastner et al., 2007; Konen and Kastner, 2008; Connolly et al., 2015).

However, phase-encoded designs provide information only about tuning location—the preferred value within a parameter space (here, preferred direction in saccade planning and execution)—and do not provide information about tuning width, that is, how narrowly or broadly a voxel responds around its preferred planned direction. Although studies using phase-encoded designs consistently report contralateral preferred tuning in human, little is known about the specificity of these responses (Sereno et al., 2001; Kastner et al., 2007; Leoné et al., 2014; Connolly et al., 2015).

Two key questions remain open. First, it is unclear whether preferred tuning width follows an orderly progression along human IPS, analogous for example to the increase in receptive-field size observed along the visual hierarchy (Harvey and Dumoulin, 2011), or whether tuning width is distributed in the absence of a clear topographic pattern. Second, the relationship between tuning width and preferred planned direction at the population level is unknown. For example, contralateral tuning in IPS could arise from (i) broadly tuned responses that remain active across much of the experimental cycle, or (ii) narrowly tuned responses that are strongly biased toward the contralateral field—or from intermediate combinations of these possibilities (Schluppeck et al., 2005).

Non-human primate neurophysiology studies of the posterior parietal sulcus predominantly focus on studying visual field position, with few investigating planned preferred saccade direction (Ben Hamed et al., 2001; Wardak et al., 2004; Patel et al., 2010; Arcaro et al., 2011). Recent studies that do report preferred saccade direction describe a contralateral organization in monkey lateral intraparietal area (Savaki et al., 2010; D’Souza et al., 2025; Griggs et al., 2025). However, these studies do not provide information about preferred tuning width.

Modern forward-modelling approaches, combined with high-field functional MRI, allow the estimation of both tuning direction and tuning width in humans, in vivo (Dumoulin and Wandell, 2008). Here, we apply this approach to saccade planning and execution using the memory-guided saccade population receptive-field approach (mgspRF). This framework enables us to derive maps of preferred planned saccade direction and their associated tuning widths across human early visual cortex and IPS. Our goal is to address open questions about tuning width in memory-guided saccades and to help bridge the gap between human functional imaging and non-human primate neurophysiology. To this end, we acquired ultra-high-field (7T) data to probe large-scale organizational principles of preferred direction and tuning width in saccade planning and execution, in humans, in vivo (Dumoulin et al., 2017a; Fabius et al., 2022). Our results show a clear contralateral organization of eye movement planning and execution and provide the novel characterization of tuning width topography in human IPS.

## Methods

### Participants

Eleven healthy human participants took part in the memory-guided saccade mapping measurement. Eighteen healthy human participants took part in the pRF mapping measurement. Each participant gave written informed consent. All experimental procedures were approved by the local ethics committee at the School of Medical, Veterinary and Life Sciences of the University of Glasgow (reference number: 200180191 and GN19NE455). Participants completed two or three scanning sessions, with one training session being completed before the first scanning session, giving a total of 26 scanning sessions and 11 training sessions.

### Projector & eye tracker

Stimuli were projected at 120 Hz with a PROPixx projector (VPixx Technologies, Saint-Bruno, QC, Canada) onto screen placed at the end of the scanner bore. The screen dimensions were the following: 320mm × 400mm. Participants viewed the screen through an angled mirror at 8.89° visual angle from 959 mm (57.0 pixels/degree) and the infrared mirror of the eye tracker. Eye-position data were acquired with an MR compatible Eyelink 1000 at 250 Hz (SR Research, Ottawa, ON, Canada). Stimuli were presented with MATLAB (The Mathworks, Natick, MA) using the Psychtoolbox and the Eyelink toolbox extensions (Cornelissen et al., 2002; Kleiner et al., 2007).

### MRI – data acquisition

MRI data was acquired on a 7T Siemens Magnetom Terra system (Siemens Healthcare, Erlangen, Germany) and a 32-channel head coil (Nova Medical Inc., Wilmington, MA, USA) at the Imaging Centre of Excellence (University of Glasgow, UK). We collected T1-weighted MP2RAGE anatomical scans for each subject (0.625 mm isotropic, FOV = 160×225×240 mm3, 256 sagittal slices, TR = 4680ms, TE = 2.09ms, TI1 = 840 ms, TI2 = 2370ms, flip angle1 = 5°, flip angle2 = 6°, bandwidth = 250Hz/px, acceleration factor = 3 in primary phase encoding direction). Total acquisition time was 12 minutes.

During the tasks, functional data were acquired as T2*-weighted echo-planar images (EPI), using the CMRR MB (multi-band) sequence with the following acquisition parameters: resolution = 1.5 mm isotropic, FOV = 192 × 192 × 84 mm^3^, repetition time (TR) = 2000 ms, echo time (TE) = 25 ms, flip angle = 72°, multiband acceleration factor = 2, phase encoding direction = anterior to posterior. For the pRF mapping task we collected 155 volumes per run (310 s), and 162 volumes (324 s) for the eye-movement tasks (clock-wise and counter-clockwise). For each MRI session, we recorded 5 volumes with the same EPI sequence parameters with the phase-encoding direction inverted (posterior to anterior; top-up EPI, to correct for susceptibility-induced distortions and facilitate co-registration with the anatomical data. We acquired the top-up EPIs at the beginning of the MRI session (between the first and the second EPI run) and at the end of the MRI session (Andersson et al., 2003; Holland et al., 2010; Fabius et al., 2022).

### MRI – EPI pre-processing and co-registration

All analyses were performed in AFNI (Saad et al., 2006), R (Team) and Python (Rossum, 1995). Functional scans were slice timing corrected using the AFNI function 3dTshift.

First, each fMRI run was denoised using the function dwidenoise from the Mrtrix3 tools (https://www.mrtrix.org/). This denoising approach is based on principal component analysis (PCA), following the apriori knowledge that the eigenspectrum (distribution of eigenvalues) of a random covariance matrix (representing pure noise) follows a known distribution. Detailed methods are described in (Veraart et al., 2016).

For each MRI session, we computed a warp field to correct for geometric distortions from the original (non-motion corrected) EPI volumes, using the function 3dQwarp: we averaged the first 5 volumes of the EPI following the acquisition of the top-up EPI, and averaged the 5 volumes of each top-up EPI run. The resulting undistorted (warped) EPI volume is the halfway warping between the two average volumes.

Motion parameters between runs in a session were estimated by aligning the EPI volumes to the first volume of the first EPI run using the function 3dvolreg. The motion estimates and warp field results were combined and applied in a single step, using the function 3dNwarpApply. The mean EPI volume (collapsing across time after unwarping and motion correction) was co-registered to the anatomy. First, we brought the anatomy and the session mean EPI volume into the same space by aligning their respective centres of mass. Next, the ‘Nudge dataset’ plugin in AFNI was used to manually provide a good starting point for the automated co-registration. This registration consisted of an affine transformation, using the local Pearson correlation as cost function in the function 3dAllineate. The individual motion-corrected runs were then de-spiked (using the function 3dDespike), scaled to obtain percentage BOLD signal change, and detrended with a 3rd order polynomial (using the function 3dDetrend). We averaged all processed EPI runs per task and participant to increase the signal to noise ratio.

EPI volumes from the moving bar condition (see *pRF mapping – moving bar* section below), as well as from clockwise and counter-clockwise eye movement conditions (see *Memory-guided saccade mapping* section below) were co-registered to the EPI volumes of the moving bar to obtain a voxel-by-voxel correspondence between tasks. The outcome of the co-registration for each participant and session was visually checked by evaluating the location of anatomical markers as gray matter/white matter (GM and WM, respectively) and GM/cerebro-spinal fluid (CSF) boundaries in the calcarine sulcus and the parietal cortex. All analyses on the functional data were performed on GM voxels in EPI space. 2mm spatial smoothing was applied to the EPI data.

### MRI – anatomy segmentation

Data processing was conducted using Freesurfer (https://surfer.nmr.mgh.harvard.edu/), AFNI/SUMA (https://afni.nimh.nih.gov/pub/dist/doc/htmldoc/index.html), and R (https://www.r-project.org/). For each subject, we first skull stripped the MP2RAGE image, applying the AFNI function 3dSkullStrip to the second inversion image. The skull stripped anatomy was processed with the recon-all Freesurfer pipeline. Freesurfer output was converted into SUMA, using the command @SUMA_Make_Spec_FS for visualisation and processing.

### Training session

Participants completed a behavioural training session before the scanning sessions to familiarize with the tasks. The training consisted of up to two runs of each task that would be performed in the scanner: (1) the moving bar, (2) clockwise memory-guided saccade mapping (3) counterclockwise memory-guided saccade mapping. During the training, participants were seated in front of a BenQ XL2411 screen (540 × 300 mm) at a distance of 600 mm (39.6 pixels/degree). Participants placed their chin on a chin-rest and eye movements were recorded with an Eyelink 1000 at 1000 Hz (SR Research, Ottawa, ON, Canada). Verbal feedback was provided to make sure participants were maintaining fixation when required and were explicitly asked to refrain from performing anticipatory saccades in the clockwise and counter-clockwise eye-movement tasks.

### pRF mapping – moving bar

Visual field mapping stimuli consisted of contrast-defined bars of cardinal and diagonal orientations, as used in previous pRF mapping studies (Dumoulin and Wandell, 2008; Harvey and Dumoulin, 2011; Fracasso et al., 2016b; Fracasso et al., 2016a; Harvey et al., 2020). Participants fixated at a central grey fixation target (r = 0.65°; Thaler et al., 2013) while a high-contrast bar with a checkerboard pattern (∼0.6° checks) swept across a uniform gray background in 20 equally spaced steps. Each step lasted 1.6 s. Bars had a width of 1.75°, exposing approximately three rows of checks. Alternating rows moved in opposite directions of each other with a speed of 0.5°s. Bars were presented with four different orientations and swept across the screen in eight different directions (two horizontal, two vertical, four diagonal). After each 1 or 2 sweeps there was a baseline period of 12 s, in which no bars were presented, and participants trained to keep stable fixation. The experiment included a total of 8 baseline periods. The total duration of the entire run was 310 s, during which 155 volumes were acquired. Each participant completed at least 5 pRF mapping runs.

### Memory-guided saccade mapping

For memory-guided saccade tasks, the required saccade target is flashed at the start of the trial, then after a delay, then fixation disappears, and participants make an eye movement to the remembered location (Willeke et al., 2019). On the other hand, in the delayed saccade task the required saccade target stays on during the delay until the fixation is extinguished acting as go signal (Basso and Wurtz, 1998; Pare and Wurtz, 2001; Wurtz et al., 2001; Wyder et al., 2003).

In this work, participants performed a memory-guided saccade task where targets appeared at successive positions in CW or CCW direction. Targets were places at the same eccentricity, separated by 36°. Each cycle lasted 64.8s (number of target locations × trial duration = 10 × 6.48 s). In a trial, participants were instructed to fixate the fixation target at the centre of the screen, while the saccade target appeared in a peripheral location at an eccentricity of 5 +/- 0.3 degrees of visual angle for 1.54s. Participants were requested to memorize the location of the saccade target peripheral location. 7 masks were then flashed, each lasting for 0.5s, during which participants kept fixation at the central fixation target. After the masks, participants were trained to perform a single saccade towards the remembered position of the saccade target, then saccade back to the centre of the screen within 1.44s after the last mask (Sereno et al., 2001; Konen and Kastner, 2008). Participants performed the task for five cycles during each scan run, resulting in scans lasting 324s (Figure 1A).

**Figure 1.**
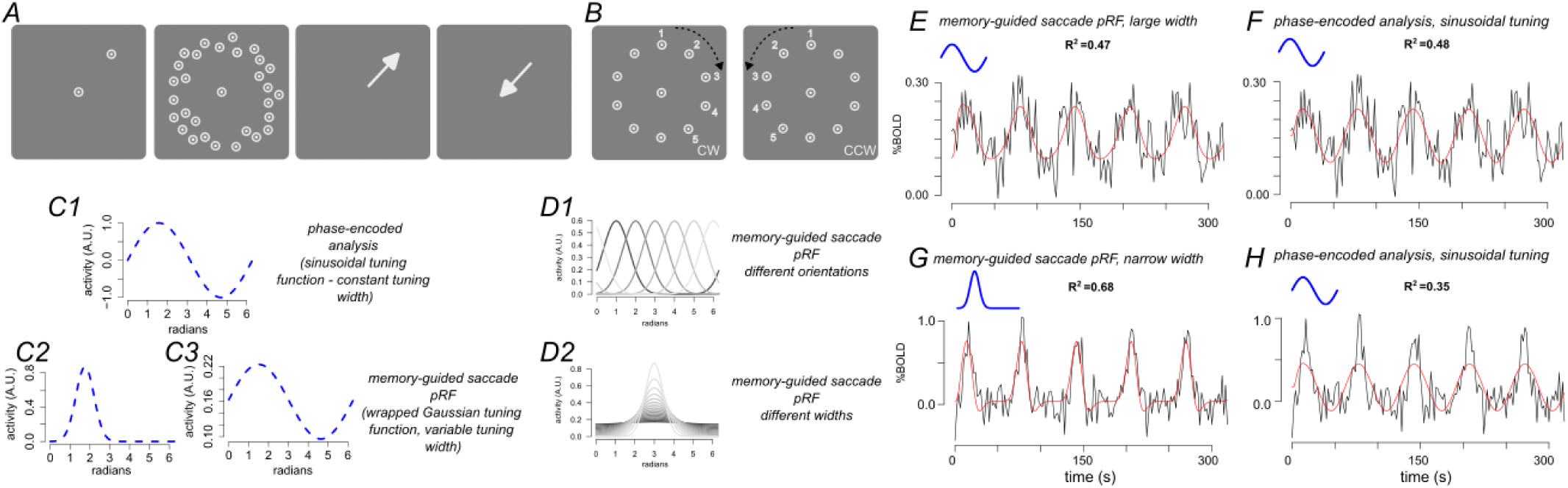
mgspRF experimental paradigm and modelling. **A.** Participants were instructed to fixate the centre of the screen, the saccade target appeared in a peripheral location at an eccentricity of 5 +/- 0.3 degrees of visual angle for 1.54s. Participants were requested to memorize the location of the saccade target peripheral location. 7 masks were then flashed, each lasting for 0.5s, during which participants kept fixation at the central fixation. After the masks, participants were trained to perform a single saccade towards the remembered position of the saccade target, then saccade back to the centre of the screen within 1.44s after the last mask. Participants performed the task for five cycles during each scan run. **B.** We alternated scans in which the target progressed in CW and CCW directions. **C1.** Example tuning function used in the classic phase-encoded analysis. In this case the tuning function is represented by a sinewave function defined by variable amplitude and phase. **C2-C3.** mgspRFs mask were defined as 1D wrapped Gaussians. Two of such wrapped Gaussians are represented to showcase different tuning width parameters. **D1-D2.** Different levels of preferred tuning location and preferred tuning width used in the mgspRF in the fitting stage. **E,F.** Example single-voxel time series (black line), fit with the mgspRF model and the phase-encoded model (red lines). The goodness-of-fit (R^2^) between the two fits is very similar (47% and 48%, respectively). Blue line represents winning tuning functions, showing very similar profiles in the two cases. **G,H.** Example single-voxel time series (black line), fit with the mgspRF model and the phase-encoded model (red lines). The goodness-of-fit (R^2^) is clearly better for the mgspRF case (68%) compared to the phase-encoded case (35%), as this particular voxel is characterized by a relatively narrow tuning, that can be captured by the mgspRF model, but not the phase-encoded model. Note however how both models yield relatively high goodness-of-fit (R^2^) values; although the phase-encoded model is outperformed by the mgspRF model, the phase-encoded model still captured a substantial portion of the variability in the signal.

We alternated scans in which the target progressed in CW and CCW directions and allowed participants to rest a few seconds between scans (Figure 1B). Each participant took part in two to three scanning sessions, each lasting ∼1.2 h.

### Memory-guided saccade population receptive fields (mgspRF)

MgspRFs of voxels in the GM mask were estimated as 1D wrapped gaussians (Figure 1C1-C3).

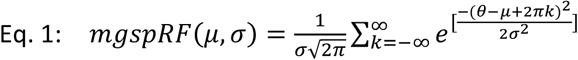

The mgspRF (Eq. 1) is defined as a function of the tuning direction (μ) and tuning width (σ), see Figure 1D1-D2.

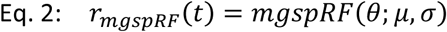

To create a predicted response (r_mgspRF_), eye movement direction over time is passed through Eq. 1, for any given pair of tuning direction (μ) and tuning width (σ). r_mgspRF_ is then temporally downsampled to match the acquisition resolution of our imaging protocol (from 6hz to 0.5hz, 2 seconds), Eq. 2.

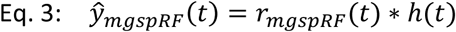

We convolved the predicted response (r_mgspRF_) with a hemodynamic response function (HRF, n1, t1, n2, t2, a2, Glover, 1999) to get a predicted BOLD response (ŷ_mgspRF_, Eq. 3). After the convolution, the predicted BOLD timeseries (ŷ_mgspRF_) were further downsampled to match the TR (0.5 Hz). We estimated the mgspRF parameters for each voxel (μ,σ) by taking the prediction that yields the maximum variance explained (R^2^). Parameters were found using an exhaustive grid search over 28512 predefined parameter combinations, for each voxel (μ=22, σ=16, n1=3, t1=3, n2=3, t2=3).

Please note that for the mgspRF we iteratively also changed the shape of the HRF used to derive the predicted response. We did so to allow HRF changes in terms of response delay as well as width, which could capture variability that might have been spuriously captured by the mgspRF tuning width. Fitting was performed using custom build code in R (https://www.r-project.org/), version 4.2.1.

### Phase-encoded analysis (travelling wave)

For each participant and voxel, we analysed the time series by fitting a sinusoid with a period of 64.8s, the same as a single revolution across 10 visual field targets. We computed the coherence between the time series and the corresponding best-fitting sinusoid and the phase of the best-fitting sinusoid (Figure 1C1). Coherence is a measure of signal-to-noise, introduced by Engel and colleagues (Engel et al., 1994; Engel et al., 1997), whose value ranges between 0 (no modulation, or small signal compared to noise) to 1 when the fMRI signal modulation at the specified frequency is large relative to the noise. Noise is defined as the total amplitude obtained by summing the signal present at the other frequency components. The resulting phase measures the temporal delay of the fMRI signal relative to the beginning of the experimental cycle and corresponds to the polar angle component of the topographic map. Different phases correspond to different target locations around the visual field (Eq. 4). c represents the coherence between the observed time series and the best fitting sinusoid. h(f,ϕ_f_) represents the sampled harmonic at the best fitting frequency and phase (ϕ_f_). On the right side of Eq. 4, the numerator, a_F_ is the amplitude of the best fitting harmonic, the denominator is the square of the summed amplitude of all the other frequency components a_f_.

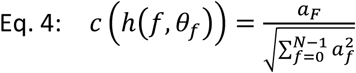

### Population receptive fields (pRFs)

PRFs of voxels in the GM mask were estimated as 2D isotropic Gaussians.

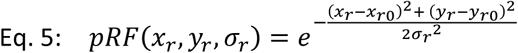

The pRF (Eq. 5) is defined as a function of retinotopic coordinates (*x_r_, y_r_*) with three parameters: horizontal center (x*_r0_*), vertical center (y*_r0_*), standard deviation (σ*_r_*). The size of the pRF is defined by σ.

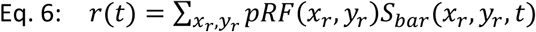

To create a predicted response (r), the pRF is multiplied elementwise with a spatially (12 pixels/°) and temporally (6 Hz) downsampled version of the moving bar stimulus (S_bar_), binarized to a contrast-defined image (Eq. 7).

We convolved the predicted response (r) for the moving bar stimulus with a standard hemodynamic response function as described above for the mgspRF to get a predicted BOLD response (ŷ, Eq. 7).

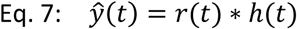

After the convolution, the predicted BOLD timeseries (ŷ) were further downsampled to match the TR (0.5 Hz). We estimated the pRF parameters for each voxel (x_r0_, y_r0_, σ) by taking the prediction that yields the maximum variance explained (R^2^). Parameters were found using an exhaustive grid search over 12100 predefined parameter combinations (22 x_r0_, 22 y_r0_, 25σ). Fitting was performed with custom build code in R (https://www.r-project.org/), version 4.2.1.

### Region of interest (ROI) definition

We obtained regions of interest (ROIs) from the Wang et al., 2015 atlas (Wang et al., 2015), which provides the location and extent of 25 topographic maps from a large population of individual subjects (N = 53) and is available in surface- and volume-based standardized space Montreal Neurological Institute’s 152 brain template (MNI152).

In the current study, we selected the following ROIs: V1 (combination of V1ventral and V2dorsal), V2 (combination of V2ventral and V2dorsal), V3 (combination of V3ventral and V3dorsal), V3A and V3B hV4, temporal occipital cortex 1 and 2 combined (hMT-MST), intraparietal sulcus areas 0 to 4 (IPS0-4 (Swisher et al., 2007). ROIs were initially defined on the surface and projected into the underlying anatomy using the function 3dSurf2Vol (Saad and Reynolds, 2012).

Please note that we report average maps across individuals only for visualization purposes. All the descriptive and statistical analyses reported in the manuscript are based on participant data in native space.

### Statistics

#### Adjusted R^2^

The mgspRF model and the phase-encoding model are characterized by a different number of parameters. To compare the goodness-of-fit between the two models it is necessary to account for the different number of parameters. In general, a model with more parameters can capture more variability than a more parsimonious model, just on the account of the number of parameters per se, artificially driving a better goodness-of-fit. To avoid bias and allowing for a fair comparison between the mgspRF model and the phase-encoding model we computed and compared the adjusted R^2^ for each model. Adjusted R^2^ penalizes R^2^ goodness-of-fit by the number of parameters used in the model, thus correcting for potential overestimations.

#### Linear mixed-effects model and t-tests

We analysed the trends along cortical hierarchy in our models a linear mixed-effects model. In these analyses, for each participant and ROI, we selected all voxels within the ROI, computed the mean of the variable of interest and took this as the estimate for the specific participant and ROI. In the model, we used the participant as a random effect component. Models were fitted using the “lme4” package in R (Bates et al., 2015a; Bates et al., 2015b). We compared post-hoc differences between models and model results between regions of interest, across participants, using Bonferroni corrected paired t-tests.

#### Receiver-operating-characteristic curve (ROC) and area-under-the-curve (AUC)

The receiver operating characteristic curve (ROC) showcases the results of a binary classification based on the true positive rate and false positive rate at each threshold setting (Hautus et al., 2021). True positive rate and false positive rate range between 0 and 1. The area-under-the-curve represents the area under the ROC curve. The closer AUC is to 1, the better the overall binary classification.

In our experiment, for each participant and ROI, we assessed saccade direction specificity across voxels comparing the saccade direction distribution between the left and the right hemispheres. We have obtained ROC and AUC for each participant and ROI, representing an index of separability between the two hemispheres. If the two distributions were overlapping (not separable), we would observe AUC values close to 0.5, indicating no difference between the left and right hemisphere, hence, the lack of contralateral specificity. On the other hand, AUC values closer to 1 indicate good separability between the hemispheres, hence, clear contralateral specificity.

In this context, AUC tells us how the classes (hemispheres) can be separated by the distribution of tuning location (preferred saccade direction).

## Results

### mgspRF model and comparison with the standard phase-encoded model (travelling wave)

Here we applied a forward modelling approach, inspired by the standard population receptive field approach (pRF), to the classic memory-guided saccade paradigm, used to identify the preferred saccade direction for memory-guided saccades (Engel et al., 1994; Sereno et al., 2001; Dumoulin and Wandell, 2008).

In this paradigm, participants make saccades to a series of memorized targets located in sequence of peripheral positions around the visual field, either in clockwise (CW) or counterclockwise direction (CCW). Saccade target targets is briefly presented, and then masked. Participants are asked to remember the target location for several seconds before performing the requested eye movement (Figure 1).

Participants performed this task while we measured blood-oxygen-level-dependent (BOLD) changes in cortical activity. We analysed the data at the individual participant level by i) performing the classic phase-encoded analysis and deriving estimates of preferred saccade direction at the individual voxel level, and ii) fitting our proposed mgspRF model, returning estimates of preferred saccade direction, as well as preferred tuning width, indicating how precisely a voxel responded to its preferred direction. We asked whether we could observe a difference in goodness-of-fit (GOF) between our proposed mgspRF model and the classic phase-encode approach, after adjusting for the difference in the number of parameters between the models.

For each participant, we averaged the time courses obtained from CW and CCW runs (at least 3 runs per condition, per participant) and fit the phase-encoded and the mgspRF models separately, for each condition (CW and CCW averaged runs). The resulting fits were then averaged between CW and CCW to give the model output.

Results of the phase-encoded model showed contralateral maps of saccade direction, with the left hemisphere encoding for rightward saccades and vice versa. This is typically observed in similar studies in humans, with different reports indicating different levels of topographic details, from a distinction in sensitivity between the leftward and rightward hemisphere (Sereno et al., 2001; Connolly et al., 2015), to more detailed topography, localizing separate cortical areas based on phase reversals at the individual participant level (Figure 2A). These differences in the level of details reported between studies might be ascribed to the relatively low signal-to-noise ratio that is generally obtained with phase-encode designs applied to eye movements (Schluppeck et al., 2005).

**Figure 2.**
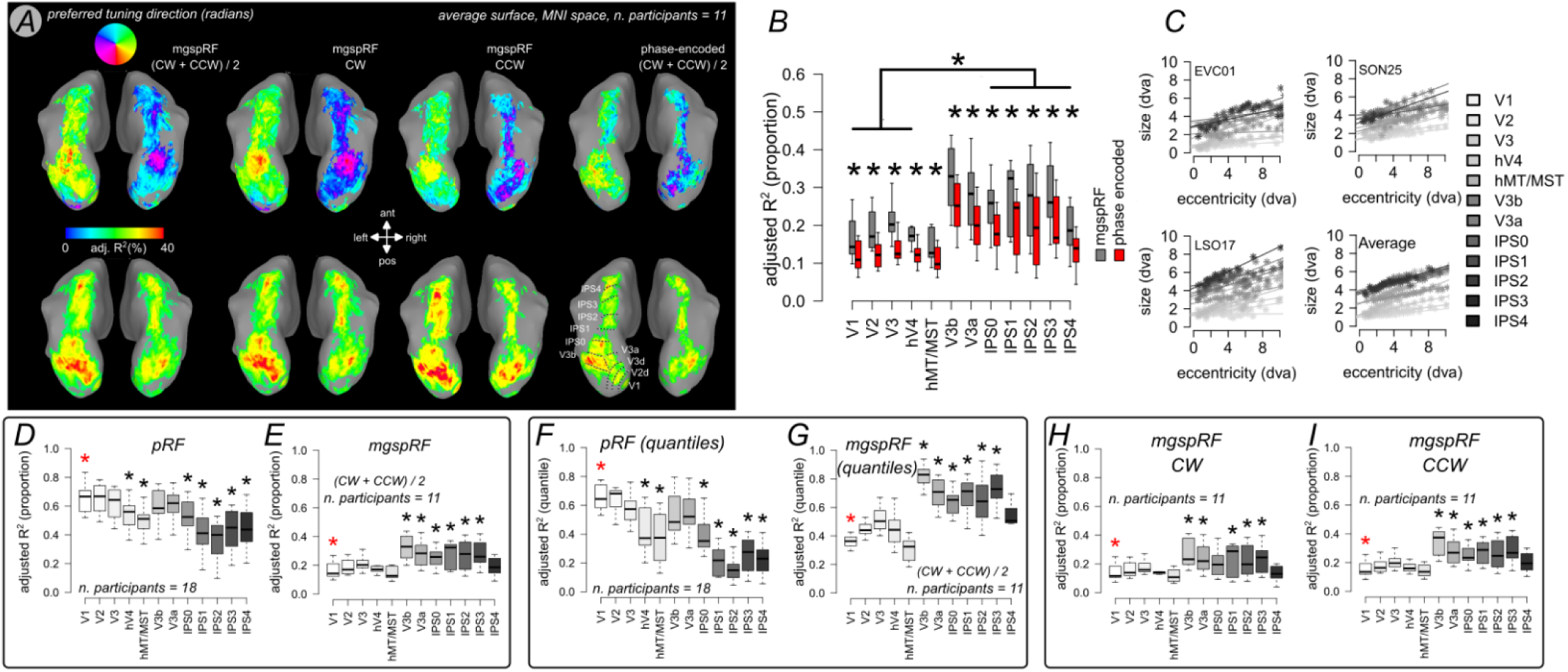
**A.** Average preferred saccade direction and adjusted variance explained maps across 11 participants from the mgspRF model and phase-encoded model. Upper line, from left to right: average fits between CW and CCW. Separate experimental condition results (CW and CCW, respectively) and results from the phase encoded model. Adjusted var. explained threshold = 15%. Both the mgspRF and the phase-encoded model showed contralateral maps of saccade direction, with the left hemisphere mapping for rightward saccades and vice versa. Please note that in the bottom right surface we present the outline of the main ROIs in the analysis. V2d and V3d refer to the dorsal portion of V2 and V3, respectively. **B.** mgspRF model captures a significant proportion of variability at the individual subject level, outperforming the adjusted goodness-of-fit estimates obtained from the phase-encoded analysis (*=p<0.001). We observe a benefit of mgspRF over phase-encoded analysis in early visual cortex (ROIs: V1-hV4), V3a/V3b, as well as in IPS (IPS0-IPS4). phase-encoded and mgspRF models yields better goodness of fit in the parietal compared to early visual cortex. **C.** Individual participant and average pRF size by eccentricity plots, showing the scaling of pRF size by eccentricity, as well as the gradual increase of average pRF size along the visual hierarchy. **D,E.** Adjusted R**^2^**across ROIs for the pRF and mgspRF, respectively. pRF fit estimates yields overall better goodness of-fit estimates compared mgspRF, with an average adjusted variance explained ∼50% for the former compared to ∼25% for the latter. In relative terms, the mgspRF yield better goodness-of-fit estimates in human IPS compared to early visual cortex. This is in stark contrast to what we observe with the standard pRF paradigm, where the trend is reversed: we see relatively better goodness-of-fit in human early visual cortex compared to IPS (black asterisks indicate significant difference with respect the reference, V1, marked with a red asterisk, *=p<0.001) **F.G.** same as D,E, but in quantiles of R^2^: it is possible to appreciate how the pRF model performs worse than mgspRF in human IPS, and vice versa for human early visual cortex. **H.I.** Adjusted R**^2^**across ROIs for the mgspRF in individual experimental conditions, CW and CCW, showing the robustness of the obtained results (black asterisks indicate significant difference with respect the reference, V1, marked with a red asterisk, *=p<0.001).

df-pRF model also showed maps of contralateral sensitivity to saccade direction between hemispheres (Figure 2A). Moreover, it captures a significant proportion of variability at the individual subject level outperforming the adjusted goodness-of-fit estimates obtained from the phase-encoded analysis. We observe a benefit of mgspRF over phase-encoded analysis in early visual cortex (ROIs: V1-hV4) as well as in IPS (IPS0-IPS4). Furthermore, phase-encoded and mgspRF yields better goodness of fit in the parietal compared to early visual cortex (Figure 2B-G).

### mgspRF tuning width changes systematically along the visual hierarchy

While the classic phase-encode model provides information about tuning location (preferred saccade direction associated with the delay period), the df-pRF model allows us to gain insights about tuning width, as well as tuning location, for each voxel. One of the advantages of in-vivo fMRI is the possibility to measure massively and in parallel across many cortical locations (voxel). The extend of the acquisition allow us to ask question about the organization (or lack thereof) of relevant parameters, especially probing the existence of topographical or columnar organization at different spatial scales. Here we test whether tuning width shows signs of topographic organization, akin to those observed in the visual domain (pRF).

pRF size is known to progressively increase along the visual hierarchy, from the small in early visual cortex to large in human IPS. The analysis of tuning width along visual hierarchy indicates a similar organization, with early visual cortex characterized by narrower tuning compared to human IPS (Figure 3). It is unlikely for this difference to be ascribed to width differences in the underlying haemodynamic response function (HRF), as during the fit stage, we allowed the shape of the HRF to change in parallel with the tuning parameters of the df-pRF model (location and width, see *Methods, Memory-guided saccade population receptive fields*). Hence, potential changes in BOLD response width could have been captured by HRF parameters.

**Figure 3.**
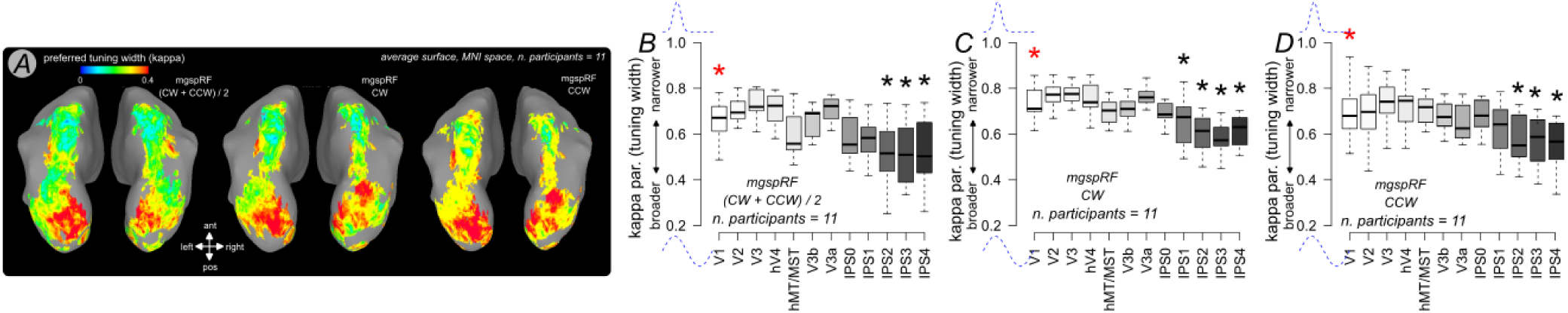
**A.** Average preferred saccade tuning width (kappa parameter). Maps are thresholded based on adjusted var. explained (threshold = 15%). The maps show a trend for narrower tuning in early visual cortex and broader tuning along the intra-parietal sulcus. **B.** Preferred saccade tuning width distribution across along the visual hierarchy for the average between CW, CCW runs. Significant differences with respect to V1 (red asterisk) are reported in black asterisks (*p<0.001). We observe significant broader tuning in the intra-parietal sulcus compared to primary visual cortex. Please note that larger values of kappa correspond to narrower tuning and vice-versa. **C,D.** Same ad panel B, here we show the preferred saccade tuning width for the CW and CCW runs, separately.

### Contralateral saccade direction specificity increases at the population level along the visual hierarchy

The mgspRF model provides us with information about saccade direction specificity (tuning width) at the single voxel level, and we have seen in the previous section how tuning width becomes broader along the visual hierarchy. However, saccade direction specificity can also change at the population level, depending on the shape of the distribution of preferred saccade direction. For example, in a given ROI, preferred saccade direction could be distributed uniformly, covering directions towards both the left and right visual fields. Alternatively, the distribution of saccade direction could be heavily skewed towards the contralateral visual field, indicating high contralateral specificity for a given ROI. These alternatives are not mutually exclusive and intermediate scenarios might also manifest.

It is possible to probe saccade direction specificity at the population level in the mgspRF model as well as the phase-encode model, across all voxels in a given ROI, by measuring the distance between the saccade direction distribution for each ROI in the left and right hemispheres. To obtain a measure of distance we measured the area under the curve (AUC) within the receiver-operating-characteristic framework in case of a binary classification (left vs right hemisphere). The closer AUC is to 1, the better the overall binary classification. In this context, AUC tells us how well the underlying classes (left and right hemispheres) can be separated by the distribution of tuning location (preferred saccade direction), per ROI and participant (Figure 4).

**Figure 4.**
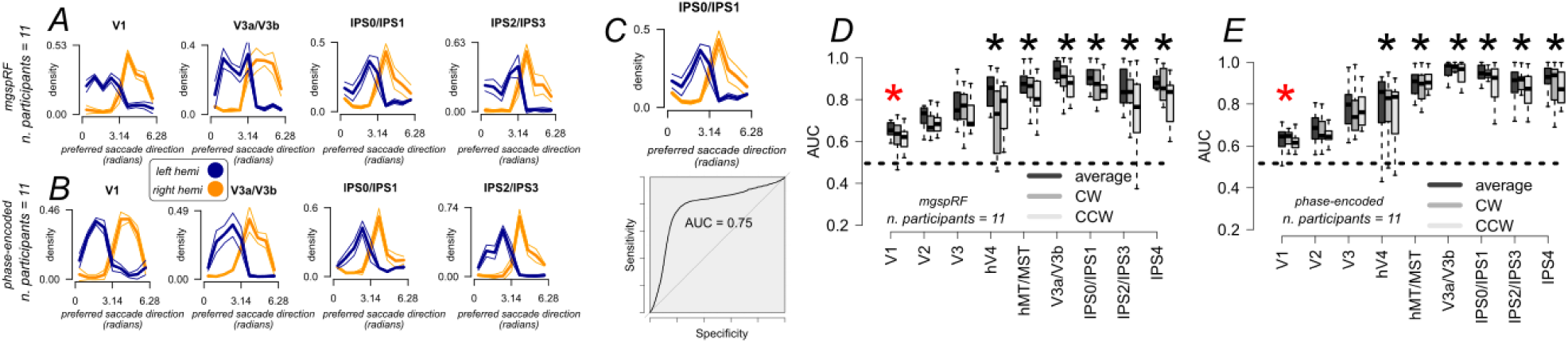
**A.** mgspRF model, average distribution of preferred saccade direction along the visual hierarchy for the left and right hemisphere, separately. Contralateral direction specificity across voxels increases at the population level along the visual hierarchy. **B.** Same as A, here we report the results of preferred saccade direction for the phase-encoded model. **C.** Example distribution for one ROI (IPS0/IPS1n in this case), and the corresponding ROC curve and AUC value, indicating the separation between the saccade tuning distributions between hemispheres. **D.** Average AUC values across the visual hierarchy, mgspRF model, for the CW and CCW average, as well as for the two experimental conditions, separately. Average AUC is significantly larger in hV4, hMT/MST, V3a/V3b and along the intra-parietal sulcus compared to primary visual cortex (*p<0.001). V1 is highlighted with a red asterisk. AUC in areas V1, V2 and V3 is significantly lower than in intra-parietal sulcus, however, preferred tuning direction in early visual cortex is significantly above 0.5 in both the phase encoded and the mgspRF model **E.** Same as D. here we report the results for the phase-encoded model.

At the population level, we observe a general trend of increasing saccade direction specificity along the visual hierarchy, with human IPS yielding higher AUC scores compared to human early visual cortex. This result is robust across the separate experimental conditions (CW and CCW) and is similar between the mgspRF model and the phase-encode model (Figure 4).

Moreover, it is important to note that while the AUC in areas V1, V2 and V3 is significantly lower than in intra-parietal sulcus, preferred tuning direction in early visual cortex is significantly above 0.5 in both the phase encoded and the mgspRF model (p < 0.01).

### mgspRF model and phase-encoded model compared to standard pRF mapping

One of the main drivers (if not the main driver) of BOLD responses in human visual cortex and IPS is visual contrast. When participants are engaged in the memory-guided saccade task, the visual input changes dramatically with every eye movement, leading to contrast changes. Given these changes, it is fair to assess whether and to what extent the classic phase-encoded design might be partially biased by contrast-based responses, as it is not clear whether spurious visual input changes could contribute the observed time series.

To answer this question, we obtained BOLD signal while participants passively viewed high contrast moving bars (4 to 6 repetitions for each participant). We used this data to fit the standard pRF model at an individual participant level. These measurements allowed us to test the eye-movement specificity of the mgspRF and phase-encoded models, compared to the visual-contrast-based specificity observed in the standard pRF paradigm.

In absolute terms, pRF fit estimates yields overall better goodness of-fit estimates compared to both mgspRF and the phase-encoded models, with an average adjusted variance explained ∼50% for the former compared to ∼25% for the latter (Figure 1C-G). This is not surprisingly as it is well-established that visual contrast is a strong driver for human early visual cortex as well as human IPS (Figure 1D-G).

However, it is interesting to note that, in relative terms, both the mgspRF and the phase-encoded models yield better goodness-of-fit estimates in human IPS compared to early visual cortex. This is in stark contrast to what we observe with the standard pRF paradigm, where the trend is reversed: we see relatively better goodness-of-fit in human early visual cortex compared to IPS. This is particularly evident when we express goodness-of-fit in terms of percentiles, rather than adjusted R^2^ (Figure 1D-G).

To perform this analysis, we converted the adjusted R^2^ obtained from every voxel for a given participant in percentiles, that is, the percentage of voxels that are below a given adjusted R^2^ value. In this way it is possible to appreciate how the pRF model performs relatively worse than mgspRF and the phase-encode model in human IPS, and vice versa for human early visual cortex. These results indicate the degree of specificity of the mgspRF and phase-encode models for the delay-saccade paradigm. Moreover, they suggest that spurious visual input changes associated with each eye movement are not the primary driver of the observed responses in the CW and CCW paradigms.

### The observed responses cannot be accounted for by contrast changes over time

To further tested whether the observed changes in BOLD signal at the individual voxel level could in principle be driven by contrast changes over time. Retinotopic receptive fields measured at fixations (pRFs) are stimulated by high-contrast stimuli systematically along each trial.

When participants move their eyes towards the periphery of the visual field and rapidly return towards the screen centre, the visual input to their eye’s changes dramatically. For example, the peripheral fixation point that participant need to memorize, the repeated high-contrast mask and the high-contrast edge at of the screen within the MRI bore. When participants move their eyes towards the remembered peripheral point, retinotopic pRFs can cross, or be partially stimulated by the screen edge All these high contrast sources can lead to potentially systematic BOLD activity (Figure 5). Thus, it is important to test whether BOLD signal at the individual voxel could in principle be accounted for by contrast changes over time (Figure 6). Here we implemented a passive (retinotopic) version of the visual stimulation, where fixation (the fovea) stayed fixed in the center, while high contrast changes occurred with respect to the static fovea, taking into account the direction and magnitude of the requested eye movement, where the screen border moves in the direction equal and opposite with respect to the saccade direction (Figure 5).

**Figure 5.**
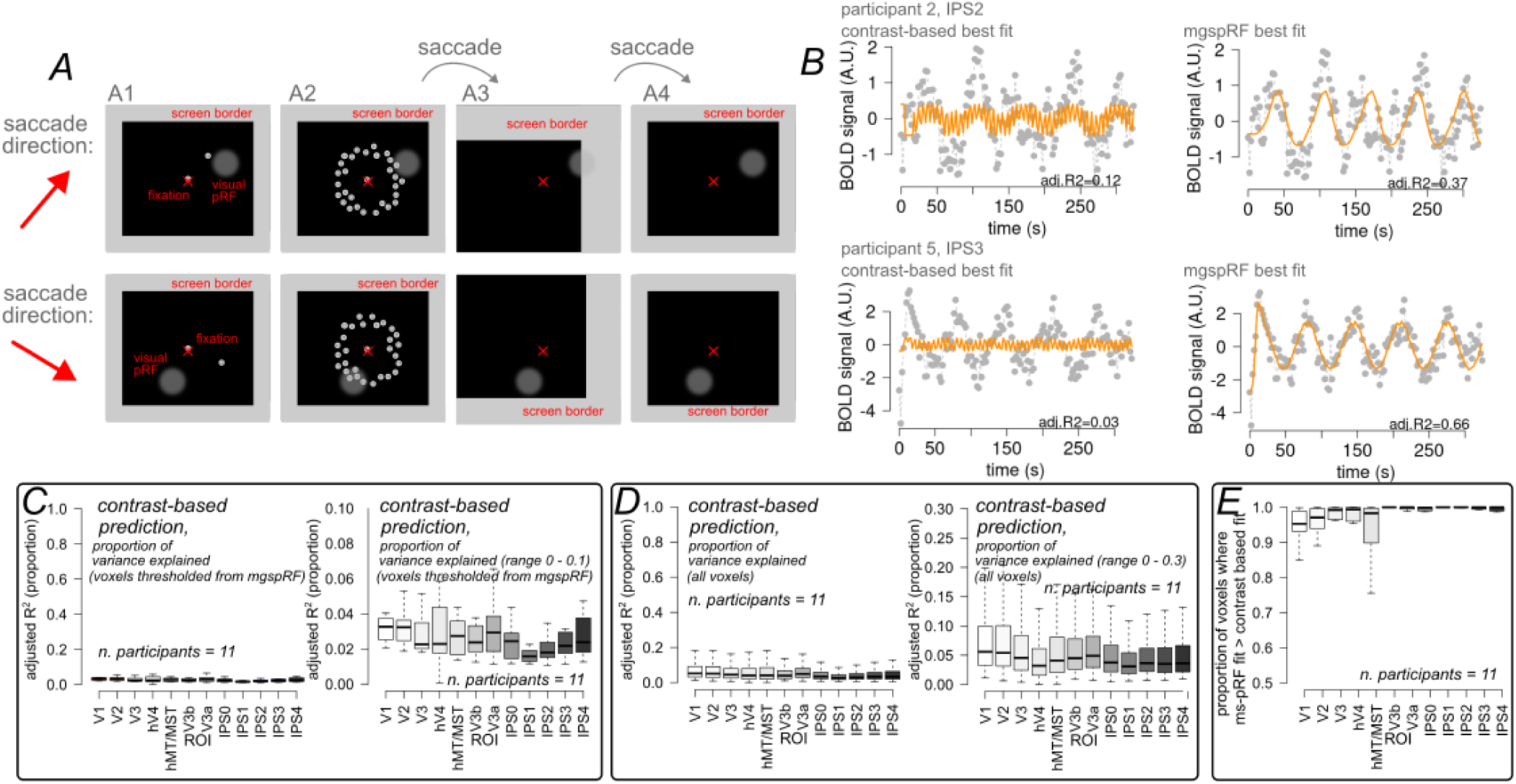
**A.** Derivation of the contrast-based predictor. Based on the visual pRF results obtained at fixation, and the stimuli sequence, we derived the predicted response based on local contrast changes. We report two examples in panel A, based on two saccade directions, where the contrast in a single pRF (grey circular area – ‘visual pRF’), changes systematically during a single trial. This change can be driven by high-contrast features appearing within the pRF (for example, due to the appearance of the multiple masks – panel A2), or because high-contrast features are brought into the pRF following a memory-guided saccade (panel A3, upper row). Here, the high-contrast feature represents the screen border within the scanner. Lateral eye movements can bring pRF closer to, or across the border, leading to systematic changes in activity. **B.** Example single voxel timecourses and the corresponding best fits for the contrast-based model as well as the mgspRF model. **C.** While some voxel show activity consistent with the contrast-based model, on average the contrast-based model accounts for less than 5% of variance explained across the visual hierarchy. Here we show the results on the same voxels that survived thresholding based on mgspRF variance explained. **D.** Same as in panel C. Here we show the results on the voxels that survived thresholding based on contrast-based prediction variance explained. **E.** Proportion of voxels where the variance explained of mgspRF model exceeded that of the contrast-based prediction, across all voxels, per ROI, without applying thresholding. More than 95% of voxels show higher variance explained for the mgspRF compared to the contrast-based prediction.

**Figure 6.**
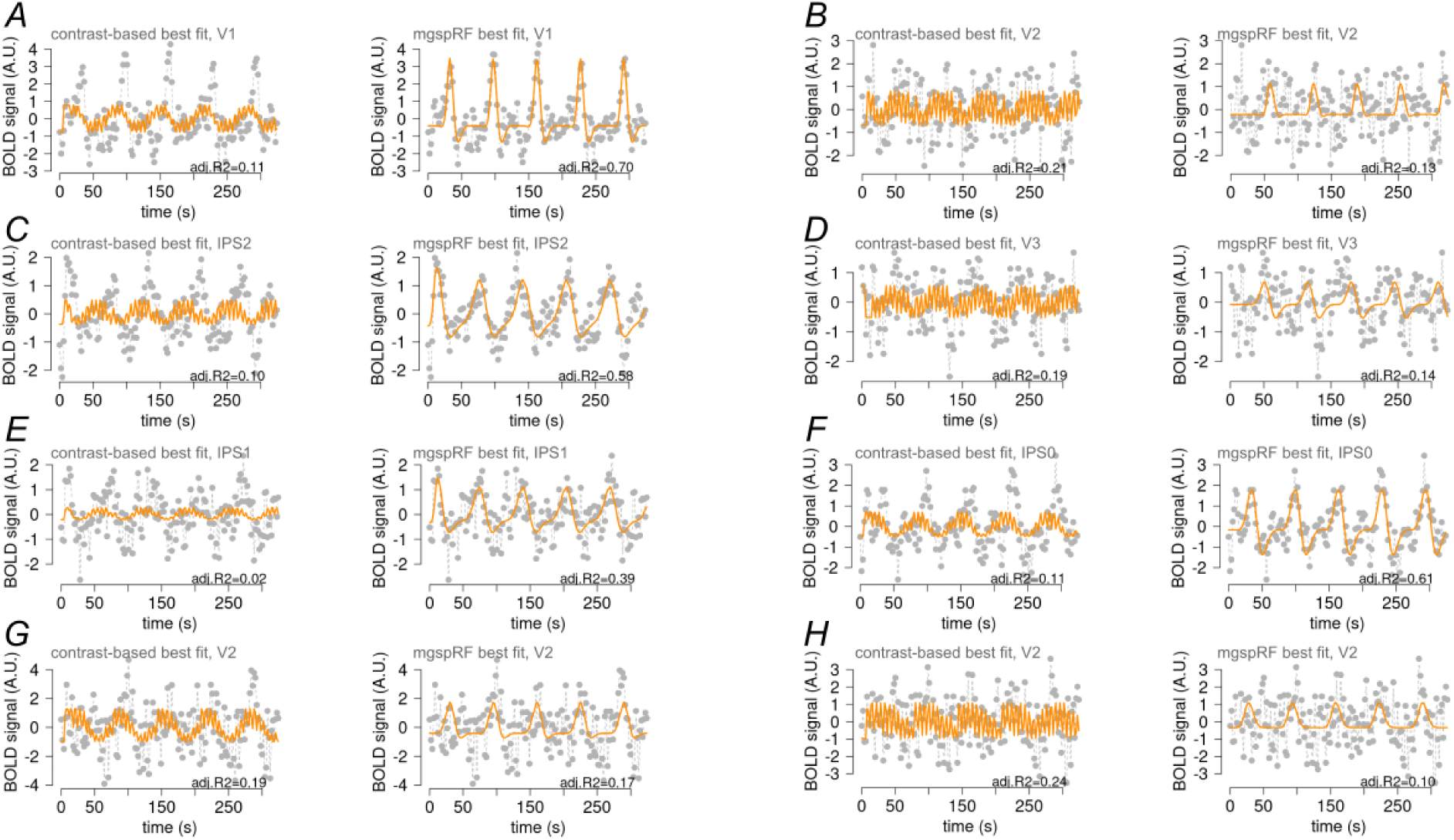
A-H. Example single voxel timecourse and the corresponding best fits for the contrast-based model as well as the mgspRF model. While more than 95% of voxels show higher variance explained for the df-PRF compared to the contrast-based prediction (see Figure 5E), there exist a small proportion of cases where the opposite is observed. Here we show a few examples, see for example panels B, G, D and H.

Using this passive (retinotopic) in combination with the visual pRFs we could obtain a predicted contrast-based time course for each individual voxel, which we tested against the observed time course using a linear model and deriving the corresponding variance explained. In general, more than 95% of voxels show higher variance explained for the df-PRF compared to the contrast-based prediction, there exist only a small proportion of cases where contras-based responses outperform the mgspRF prediction.

To further compare the goodness of fit (variance explained) of mgspRF and contrast-based models we show the proportion of voxels per each ROI exceeding a given variance explained value (namely, 10%, 15% and 25%), without thresholding (Table 1). Overall, the results indicate how mgspRF systematically outperforms the contrast-based model. Moreover, it is important to note how the mgspRF performs in early visual cortex (V1, V2, V3), showing a considerable proportion of voxels systematically exceeding the goodness of fit threshold. This result is compatible with the tuning direction results in early visual cortex, showing higher than chance preferred tuning direction.

**Table 1.** Here we report the average (and sd) proportion of voxels where the variance explained exceeded 10%, 15% and 25% for the mgspRF and the contrast-based model. There exist a significant proportion of such responses for the mgspRF model along the visual hierarchy and the intra-parietal sulcus. Few cases can be observed for the contrast-based model. To allow for an unbiased comparison, these results are based on unthresholded data, for each ROI.

| ROI: | P(voxel with<br>adj.var exp)<br>> 10% | P(voxels<br>with adj.var<br>exp) > 10% | P(voxel with<br>adj.var exp)<br>> 15% | P(voxel with<br>adj.var exp)<br>> 15% | P(voxel with<br>adj.var exp)<br>> 25% | P(voxel with<br>adj.var exp)<br>> 25% |
| --- | --- | --- | --- | --- | --- | --- |
|  | *mgspRF, mean*<br>*(sd)* | *contrast-*<br>*based, mean*<br>*(sd)* | *mgspRF,*<br>*mean (sd)* | *contrast-*<br>*based, mean*<br>*(sd)* | *mgspRF,*<br>*mean (sd)* | *contrast-*<br>*based, mean*<br>*(sd)* |
| V1 | 0.49 (0.16) | 0.10 (0.06) | 0.28 (0.14) | 0.05 (0.04) | 0.09 (0.08) | 0.01 (0.01) |
| V2 | 0.52 (0.12) | 0.09 (0.07) | 0.33 (0.11) | 0.05 (0.04) | 0.12 (0.07) | 0.01 (0.01) |
| V3 | 0.53 (0.09) | 0.07 (0.06) | 0.36 (0.08) | 0.03 (0.03) | 0.17 (0.07) | <0.01 |
| IPS0 | 0.66 (0.09) | 0.05 (0.05) | 0.48 (0.12) | 0.02 (0.02) | 0.25 (0.11) | <0.01 |
| IPS1 | 0.70 (0.12) | 0.03 (0.05) | 0.53 (0.14) | 0.01 (0.03) | 0.29 (0.15) | <0.01 |
| IPS2 | 0.68 (0.18) | 0.03 (0.03) | 0.52 (0.20) | 0.01 (0.01) | 0.28 (0.19) | <0.01 |

## Discussion

Here, we estimated voxel’s preferred tuning direction and tuning width using a forward-modeling approach (mgspRF). We demonstrate that mgspRF outperforms the classic phase-encoded approach. Importantly, mgspRF reveals a systematic increase in saccade-tuning width from early visual cortex to human IPS, uncovering a previously unknown organizational principle along human IPS. At the population level, we also show that the specificity of contralateral saccade-direction increases along the cortical hierarchy. Lastly, our measurements indicate how single-voxel responses cannot be fully explained by simple contrast-change dynamics over time.

### Preferred tuning width changes systematically along the visual hierarchy

While the classic phase-encode model provides information about tuning location (preferred saccade direction in saccade planning and execution), the mgspRF model allows us to estimate both preferred tuning location and width for each voxel. A key advantage of in-vivo fMRI is the ability to measure neural responses simultaneously across large portions of cortex, enabling us to examine how neural parameters are organized—or not organized—across cortical space. This makes it possible to test for topographic arrangement at multiple spatial scales. Here, we specifically ask whether tuning width exhibits a topographic organization, for example analogous to that observed for pRF size in the visual system.

PRF size is known to increase systematically along the visual hierarchy, from small receptive fields in early visual cortex to much larger ones in human IPS (Dumoulin and Wandell, 2008; Harvey and Dumoulin, 2011; Benson et al., 2018). Our analysis of tuning width reveals a similar pattern: early visual cortex shows narrower tuning compared to human IPS. This difference is unlikely to reflect variations in the underlying haemodynamic response function (HRF), as our fitting procedure allowed HRF shape to vary jointly with mgspRF parameters (location and width). Systematic broadening of the BOLD response due to HRF changes would therefore have been likely captured by the HRF parameters.

We observe the same pattern of results when modelling CW and CCW runs separately (split-test), demonstrating the robustness of the reported effect.

### Contralateral saccade direction specificity increases at the population level along the visual hierarchy

Preferred saccade-direction tuning width becomes broader—and therefore less specific—along the visual hierarchy at the single-voxel level. However, direction specificity at the population level depends on how preferred saccade directions are distributed across voxels within a region.

For instance, a given cortical area may show a uniform distribution of preferred directions, spanning both leftward and rightward saccades. Alternatively, the distribution may be strongly biased toward the contralateral visual field, indicating high contralateral specificity. These scenarios are not mutually exclusive.

To quantify population-level specificity, we examined the distribution of preferred saccade directions across voxels within each ROI for both the mgspRF and phase-encode models. We assessed the distance between the left- and right-hemisphere distributions by computing the area under the curve in a receiver-operating-characteristic framework.

Population-level saccade specificity increases along the visual hierarchy: human IPS shows higher AUC values—indicating better discrimination between left- and right-hemisphere direction preferences—than early visual cortex. This pattern is robust across experimental conditions (CW and CCW) and consistent across both modelling approaches. Together, these results show that contralateral specificity strengthens along the hierarchy at the population level, despite broader tuning at the single-voxel level.

Consequently, a higher-order decoder could reliably distinguish leftward from rightward saccades based on population activity in human intraparietal sulcus, even though individual voxels exhibit less precise tuning than those in early visual cortex.

It is important to note that also responses in early visual cortex show a degree of directional specificity at the population level. Furthermore, these responses in early visual cortex cannot be fully explained by contrast changes over time. This pattern suggests that the observed sensitivity to saccade planning and execution in early visual cortex is unlikely to be a spurious consequence of contrast-related confounds. Instead, it may reflect genuine sensitivity in saccade planning, consistent with recent evidence showing that early visual cortex in human and non-human primates, although generally dominated by contrast-based responses, are also modulated by effector position and direction (Weyand and Malpeli, 1993; Trotter and Celebrini, 1999; Merriam et al., 2013; Morris and Krekelberg, 2019; Fabius et al., 2022).

### Comparing the mgspRF and the classic phase-encode model

The mgspRF model provides better goodness-of-fit estimates compared to the classic phase-encode model, even after accounting for differences in model complexity. This suggests that mgspRF captures aspects of the observed data that extend beyond what is typically characterized by phase-encoding approaches (Engel et al., 1994; Engel et al., 1997; Sereno et al., 2001).

Due to the sluggishness of the haemodynamic response function (Miezin et al., 2000), it is important to note that the memory-guided saccade paradigm within the context of fMRI acquisition does not allow to disentangle the separate contributions of saccade target selection, saccade planning or saccade execution. Moreover, there is still debate as to whether the lateral intraparietal sulcus in non-human primates acts as an intention map, or selects the environmental targets that will be further processed (Andersen, 1995; Goldberg et al., 2006; Andersen and Cui, 2009; Bisley and Goldberg, 2010). The current work does not allow to disentangle between these different scenarios. Here we provide novel insights about the organizational properties of human IPS, both globally and at the individual voxel level, in a task involving saccade planning *and* execution.

Both the mgspRF and phase-encode models revealed contralateral maps of saccade direction, with the left hemisphere preferring rightward saccades and the right hemisphere preferring leftward saccades. This contralateral organization is a well-established finding in the literature, although studies differ in the level of topographic detail they report. Prior work ranges from broad hemispheric distinctions in directional sensitivity (Connolly et al., 2015) to more fine-grained topographies that identify distinct cortical areas based on phase reversals at the individual-participant level (Schluppeck et al., 2005; Levy et al., 2007). While the overall contralateral pattern of saccade specificity is consistently observed, the precise boundaries between neighbouring cortical areas—and the degree to which they can be resolved in individual participants—vary across studies.

Differences in the level of topographic detail reported across studies may stem from the relatively low signal-to-noise ratio typically associated with memory-guided saccade paradigms. By contrast, block designs in which participants alternate between rapid eye movements and fixation generally yield higher signal-to-noise ratios (Schluppeck et al., 2005; Kastner et al., 2007), however, block designs do not allow estimation of saccade-direction preferences. Studies using the memory-guided saccade paradigm vary in the number of fMRI runs (CW and CCW) collected per participant, ranging from as few as 3–4 to more than 15 (Sereno et al., 2001; Schluppeck et al., 2005). This variation directly affects the amount of signal available and, consequently, the level of topographic detail that can be resolved.

The optimal number of experimental runs depends on the goals and scope of the specific experiment, as well as on the imaging hardware. Mapping fine-grained cortical boundaries at the individual-participant level typically requires more repetitions than testing large-scale organizational principles. Higher field strengths (e.g., 7T) also provide improved signal-to-noise ratios compared to 3T scanners. In the present study, we used high-field (7T) data to investigate large-scale organizational principles of preferred saccade direction and, in particular, tuning width—measured in vivo in humans for the first time.

### Memory-guided saccade responses and visual pRFs

The mgspRF and the phase-encoded models yield better goodness-of-fit estimates in human IPS compared to early visual cortex. Results with the standard pRF paradigm show the opposite trend: relatively better goodness-of-fit in human early visual cortex compared to IPS. This result is not surprisingly as it is well-established that visual contrast is a strong driver of human early visual cortex as well as human IPS.

In the CW and CCW memory-guided saccade paradigm participant regularly move their eyes towards the edge of the field of view and back towards the centre. With every eye movement the visual input changes suddenly, thus leading to a potential confound. If responses in the CW/CCW paradigm were primarily driven by changes in contrast-based responses, we would expect to observe the same goodness of fit trend along the visual hierarchy we observe for pRF modelling, however the opposite trend is observed.

Furthermore, the goodness-of-fit pattern between early visual cortex and human IPS is unlikely explained by systematic changes in visual contrast over time. A model explicitly incorporating individual visual pRFs and contrast changes based fails to account for variance at a level comparable to the mgspRF model. It is important to note that the contrast-based model yielded relatively low goodness of fit values in early visual cortex. This is not surprising, as the memory-guided saccade paradigm involves strong visual contrast changes only in the presence of the visual masks presentation. These masks are presented at a relatively fast pace (one mask every 0.5 seconds). Such high pace of visual stimulation cannot easily be captured by corresponding changes in %BOLD signal change, due to the slow temporal features of the human haemodynamic response function (Miezin et al., 2000). This would in turn lead only to relatively small predicted variations in %BOLD signal, as can be seen in the high frequency component of the contrast-based model time series (see Figure 5 and 6). A slower component of the contrast-based responses is driven by the interaction between the visual pRF and the high contrast border of the screen within the scanner bore (see Figure 5A). While this component can capture the temporal features present in some voxels (see time series reported in Figure 5 and 6), it does not dominate the responses in the contrast-based model. Having said that, it is important to note how the contrast-based model can successfully capture a reasonable fraction of variance in more than 10% of voxels in human early visual cortex.

### Mapping saccade behaviour in human intra-parietal sulcus

We show how preferred saccade tuning width derived from a memory-guided saccade task is arranged along an orderly progression between human early and intra-parietal sulcus, following a general principle of cortical organization seen, for example, within the domain of vision, where receptive field increases along the visual hierarchy along the dorsal stream (Harvey and Dumoulin, 2011).

The contralateral sensitivity generally observed using phase-encoded designs in human intra-parietal sulcus can likely be ascribed to the presence of broad and contralaterally tuned responses. These results complement recent evidence from macaque neurophysiology in the posterior parietal sulcus, describing subregions tuned to different directions with a rough topography along the anterior-posterior axis of macaque LIP (Griggs et al., 2025).

Importantly, our results indicate how responses in early visual cortex show a degree of sensitivity for the memory-guided saccade paradigm. Interestingly, responses in early visual cortex cannot be fully accounted for by passive contrast changes over time, and show a small, albeit significant contralateral sensitivity. This is compatible with studies highlighting how early visual cortex, is also sensitive to other sensory domains beyond vision, for example encoding eye position information (Weyand and Malpeli, 1993; Trotter and Celebrini, 1999; Merriam et al., 2013; Morris and Krekelberg, 2019; Fabius et al., 2022).

## Conclusion

We show how preferred tuning width derived from a memory-guided saccade task is arranged along an orderly progression between human early and intra-parietal sulcus. Moreover, our results show how early visual cortex, while dominated by contrast-based responses, is also sensitive to saccade planning and execution. Our results fill a gap in our understanding of the functional organization of human intra-parietal sulcus by providing evidence about functional organization and topography of tuning direction and width along the visual hierarchy during saccade planning and execution.

## Acknowledgements

A.F. was supported by a grant from the Biotechnology and Biology Research Council (BBSRC, grant number: BB/S006605/1) and the Bial Foundation (Bial Foundation Grants Programme; Grant id: A-29315, number: 203/2020, grant edition: G-15516). A.B. was supported by a grant from the Italian Minister of Research and University (PRIN 2022_PNRR-P2022ST78T).

